# SCP Viz – A universal graphical user interface for single protein analysis in single cell proteomics datasets

**DOI:** 10.1101/2023.08.29.555397

**Authors:** Ahmed Warshanna, Benjamin C. Orsburn

## Abstract

Single cell proteomics (SCP) requires the analysis of dozens to thousands of single human cells to draw biological conclusions. However, assessing of the abundance of single proteins in output data presents a considerable challenge, and no simple universal solutions currently exist. To address this, we developed SCP Viz, a statistical package with a graphical user interface that can handle small and large scale SCP output from any instrument or data processing software. In this software, the abundance of individual proteins can be plotted in a variety of ways, using either unadjusted or normalized outputs. These outputs can also be transformed or imputed within the software. SCP Viz offers a variety of plotting options which can help identify significantly altered proteins between groups, both before and after quantitative transformations. Upon the discovery of subpopulations of single cells, users can easily regroup the cells of interest using straightforward text-based filters. When used in this way, SCP Viz allows users to visualize proteomic heterogeneity at the level of individual proteins, cells, or identified subcellular populations. SCP Viz is compatible with output files from MaxQuant, FragPipe, SpectroNaut, and Proteome Discoverer, and should work equally well with other formats. SCP Viz is publicly available at https://github.com/orsburn/SCPViz. For demonstrations, users can download our test data from GitHub and use an online version that accepts user input for analysis at https://orsburnlab.shinyapps.io/SCPViz/.

**Abstract graphic:** 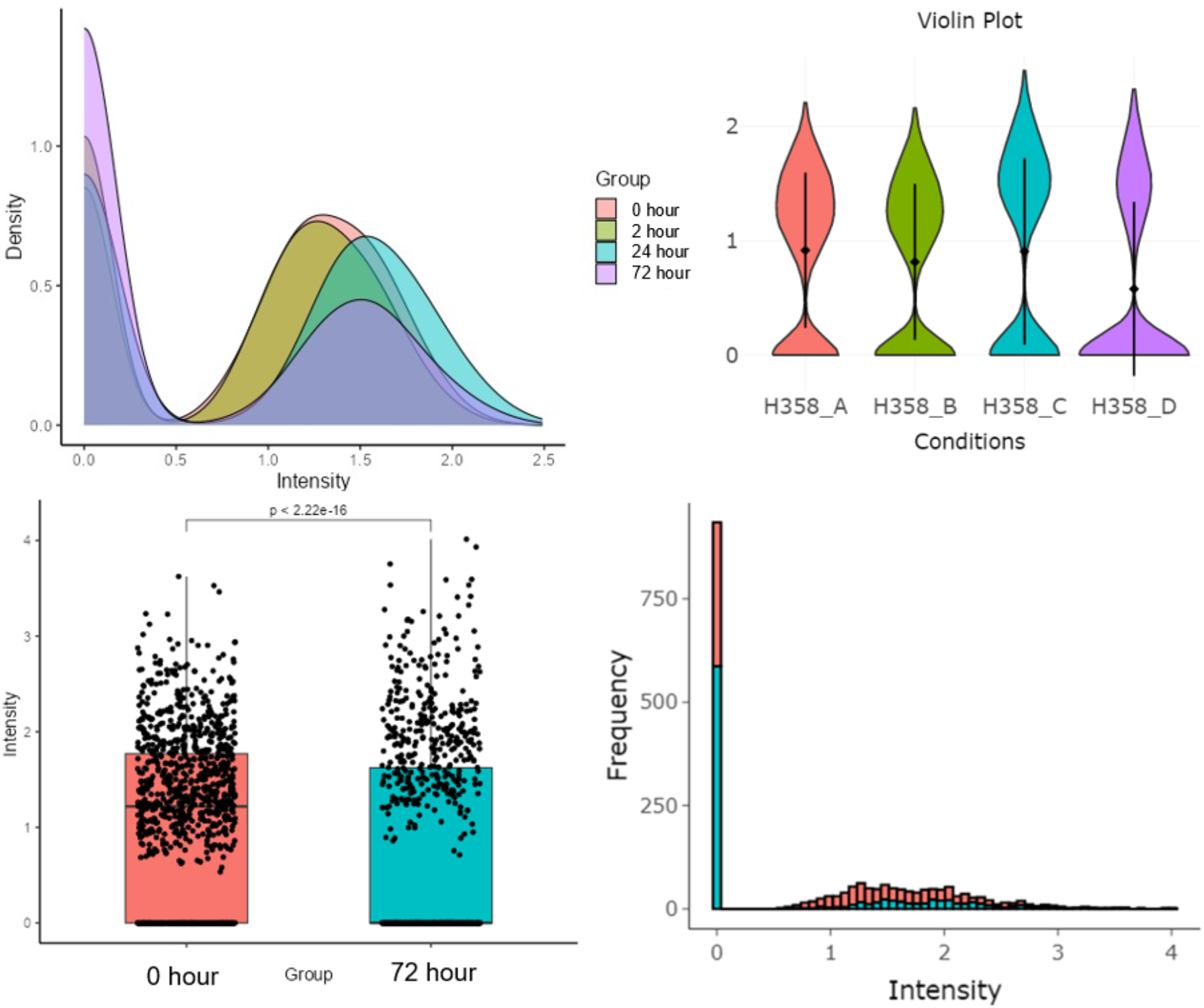

## Introduction

Single cell proteomics (SCP) by LCMS is an emerging field of research today, driven by advancements in all aspects from sample prep to instrumentation and informatics.^1,2^ Given the variety of instruments and processing tools available, performing downstream analysis, especially when evaluating the abundance of individual proteins across large datasets, becomes particularly challenging. It is not surprising, considering that proteomics as a field generally tends to lack downstream interpretation and visualization tools.^3,4^ The tools that do exist for SCP primarily focus on identification of patterns and identifying subpopulations of cells exhibiting shared phenotypes. Many of these tools employ linear or nonlinear data reduction methods for visualization, which, in nearly all cases, results in alteration or loss of the original data.^5,6^

Today, numerous packages exist for the downstream statistical analysis of proteomics data, including LFQAnalyst,^7^ MSStats,^8,9^ StatsPro,^10^ EatOmics,^11^ Amica,^12^ ProVision,^13^ EinProt^14^ and Perseus^15^. Of these, only Perseus has demonstrated the capability to visualize data across hundreds of individual cells.^16^ The SCP field continues to advanced rapidly, as demonstrated by new fit-for-purpose informatics such as IceR,^17^ PepDesc,^18^ and the R/Bioconductor toolkit “scp”^2,5^. These tools enhance our depth of coverage and improve our interpretation of these new data types. However, it could be argued that these solutions, being bioinformatics packages, are tailored for informaticians, making them less accessible to the broader proteomics or biology community.

To address these shortcomings in our research, we set out to create a toolkit compatible with every tool used in our lab. We aimed for compatibility with today’s most popular search tools, including SpectroNaut,^19^ DIA-NN,^20^ FragPipe,^21,22^ Proteome Discoverer,^23^ and MaxQuant.^24^ Additionally, we sought the ability to rapidly determine whether a protein changed between groups. The ultimate goal of SCP Viz was to adapt to new insights, such as the discovery of subpopulations of cells based on other phenotypic data.

## Methods

### SCP Viz Dependencies

SCP Viz 1.0.0 was developed using base R (R version 4.2.2 (2022-10-31)) in the MacOS operating environment with visualization provided through the Shiny application framework. The dependencies and their respective versions are as follows: “shinythemes_1.2.0”, “janitor_2.2.0”, “DT_0.28”, “gridExtra_2.3”, “tidyr_1.3.0”, “dplyr_1.1.2”, “readr_2.1.4”, “ggpubr_0.6.0”, “readxl_1.4.2”, “factoextra_1.0.7”, “FactoMineR_2.8”, “plotly_4.10.2”, “ggplot2_3.4.2”, and “shiny_1.7.4.1”.

### TIMSTOF Multiplexed proteomic data used for script construction

Work in our lab recently investigated pancreatic cancer cell lines treated with a KRAS^G12D^ inhibitor.^25^ We manually extracted the Proteome Discoverer output data for PANC 08.13 cells, which were treated with the inhibitor for 48 hours, into a CSV file for the generation of the SCP Viz tool kit. To simplify visualization, we removed the blank and carrier channels from the analysis. Furthermore, because of distortions caused by isotopic impurities in the residual carrier, we excluded single cells labeled with 134 from the analysis. The resulting output file contained data from approximately 560 single cells.

### Reanalysis of Vegvari *et al*., single cell chemical proteomics data

The processed results from Vegvari *et al*.,^26^ were downloaded from the PRIDE repository (PXD025481). These were converted to CSV files in Excel, where we replaced “:” in the sample headers with “_” using Excels’ Find/Replace function. Although the Shiny package could visualize the complete files, the abundance in the carrier channel compressed the scale, complicating interpretation. The carrier channels labeled with TMT 131 were manually deleted to generate the figures used in this manuscript. The resulting CSVs were uploaded to SCP Viz and conducted an analysis under two conditions: control and treated. The TOP1 and TYMS were chosen as proteins for analysis due to their significance in the original study.

### Importing SpectroNaut data from single human PANC 02.03 cells

Single cell data previously reported by our group that was generated on a TIMSTOF Single Cell Proteomics system^27^ was exported from SpectroNaut 18 in the default .TSV report template. The report was opened in Excel and all periods in the header column were replaced with underscores using the Find and Replace function. The resulting output report was saved as CSV. The data was imported into SCP Viz using “PG_Quantity” as text used for the quan columns.

## Results

### SCP Viz can import single cell data in all normal proteomics formats

Currently, there are over 1,000 data processing tools for shotgun proteomics.^28^ While this number likely does not reflect currently supported pipelines, it underscores the challenge of developing universal downstream tools for interpretation. SCP Viz addresses these challenges by using combinatorial text filters. A summary of the input text strings required for SCP Viz is provided in **Table 1**.

**Table 1.**
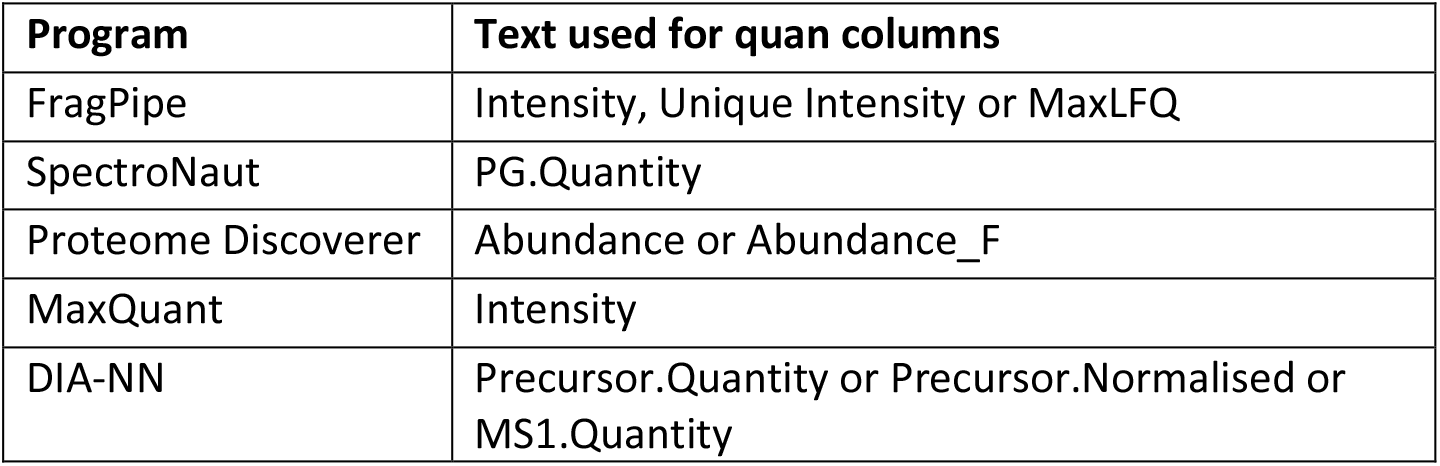
A summary of the text filters used to identify single cell quantification values in some popular proteomics programs.

| Program | Text used for quan columns |
| --- | --- |
| FragPipe | Intensity, Unique Intensity or MaxLFQ |
| SpectroNaut | PG.Quantity |
| Proteome Discoverer | Abundance or Abundance_F |
| MaxQuant | Intensity |
| DIA-NN | Precursor.Quantity or Precursor.Normalised or MS1.Quantity |

### SCP Viz simplifies visualization of data generating using TIMSTOF based SCP

As previously stated, visualizing SCP data can be a considerable challenge, particularly at the level of individual proteins. It is challenging to illustrate how visualizing protein expression across hundreds or thousands of cells using traditional proteomics tools, such as Proteome Discoverer, can be time consuming in their current software versions. SCP Viz, however, relies on text documents and streamlined integrated statistics in R, facilitating near instantaneous visualization of data across various plots. As one example, our recent research demonstrated that the KRAS^G12D^ inhibitor, MRTX1133, results in the downregulation of the MAPK pathway in mutant pancreatic cancer cells lines. **Figure 1** is a representation of MAPK1 expression in approximately 250 control and treated cells using plots toggled in SCP Viz. From the recapitulated mean and standard deviation of the data alone (**Figure 1A**) it is difficult to assess the impact of drug treatment on these cells. However, switching to a box plot with integrated statistics reveals a significant alteration, despite the relatively small alteration in the mean protein abundance (**Figure 1B**). A density plot provides further insight into protein expression with the majority of treated cells appearing to congregate in the area of lowest relative expression. (**Figure 1C**)

**Figure 1.**
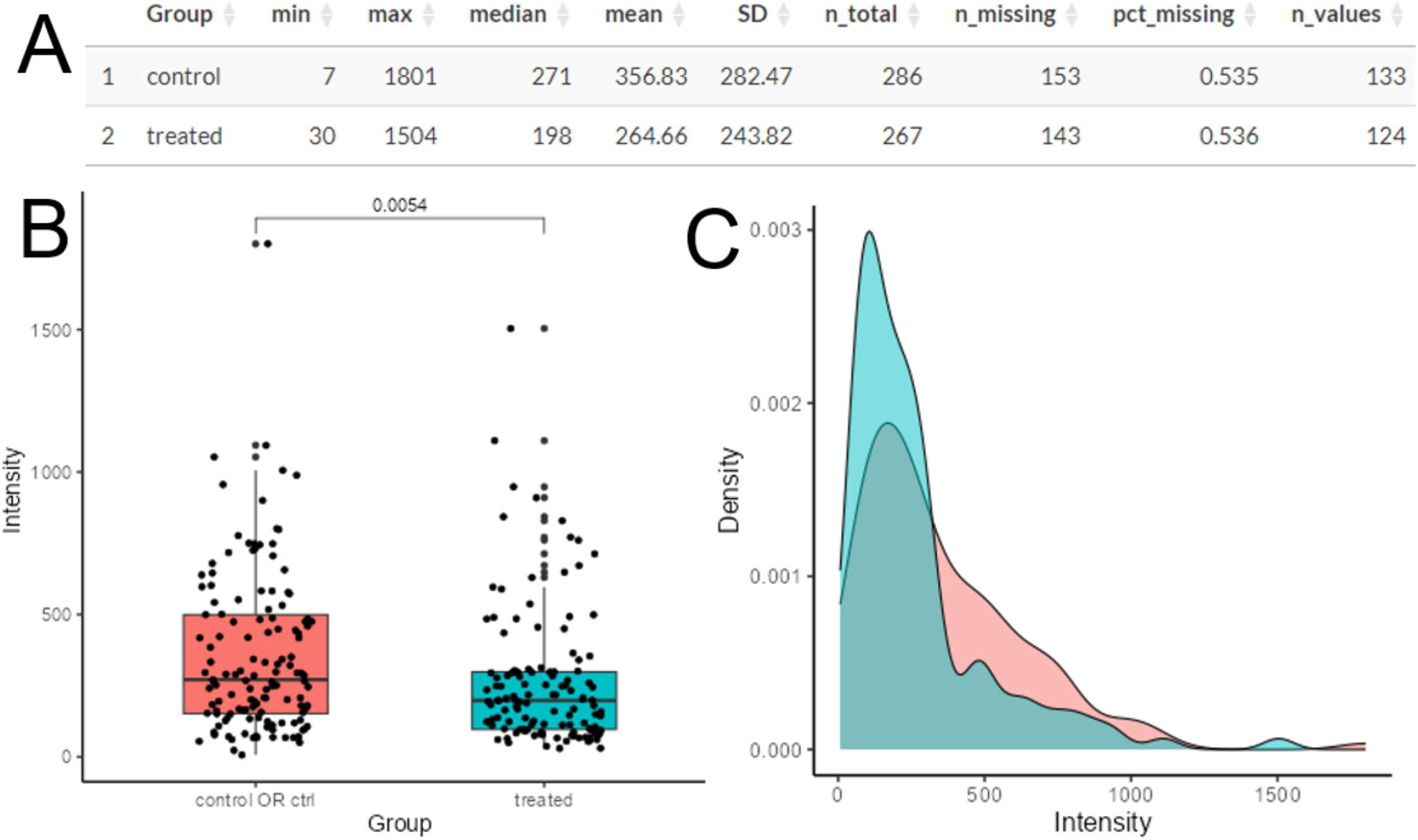
An analysis of MAPK1 expression in cells treated with a KRAS inhibitor. **A**. A summary table of the study displays the mean, median, standard deviation and number of missing values for this protein across all analyzed cells. **B**. A boxplot showcasing the mean, standard deviation and p-value visualized. **C**. A density plot of illustrating MAPK1 protein expression across all single cells.

### SCP Viz can reproduce previous findings using Orbitrap based SCP

In a recent study, Vegvari *et al*. performed a chemoproteomic analysis of A549 cancer cells treated with chemotherapeutics in an evaluation of single cell commitment to death.^26^ The processed results from this study were downloaded from the PRIDE repository and were re-evaluated with SCP Viz. A notable finding of the original study was the differential expression of TOP1 after 48 hours of treatment with campothecin. As shown in **Figure 2**, SCP Viz identifies a significant alteration in TOP1 expression. However, given the number of inputs, plots of the original unadjusted expression appear to provide a clearer visualization of the cellular heterogeneity than box plots.

**Figure 2.**
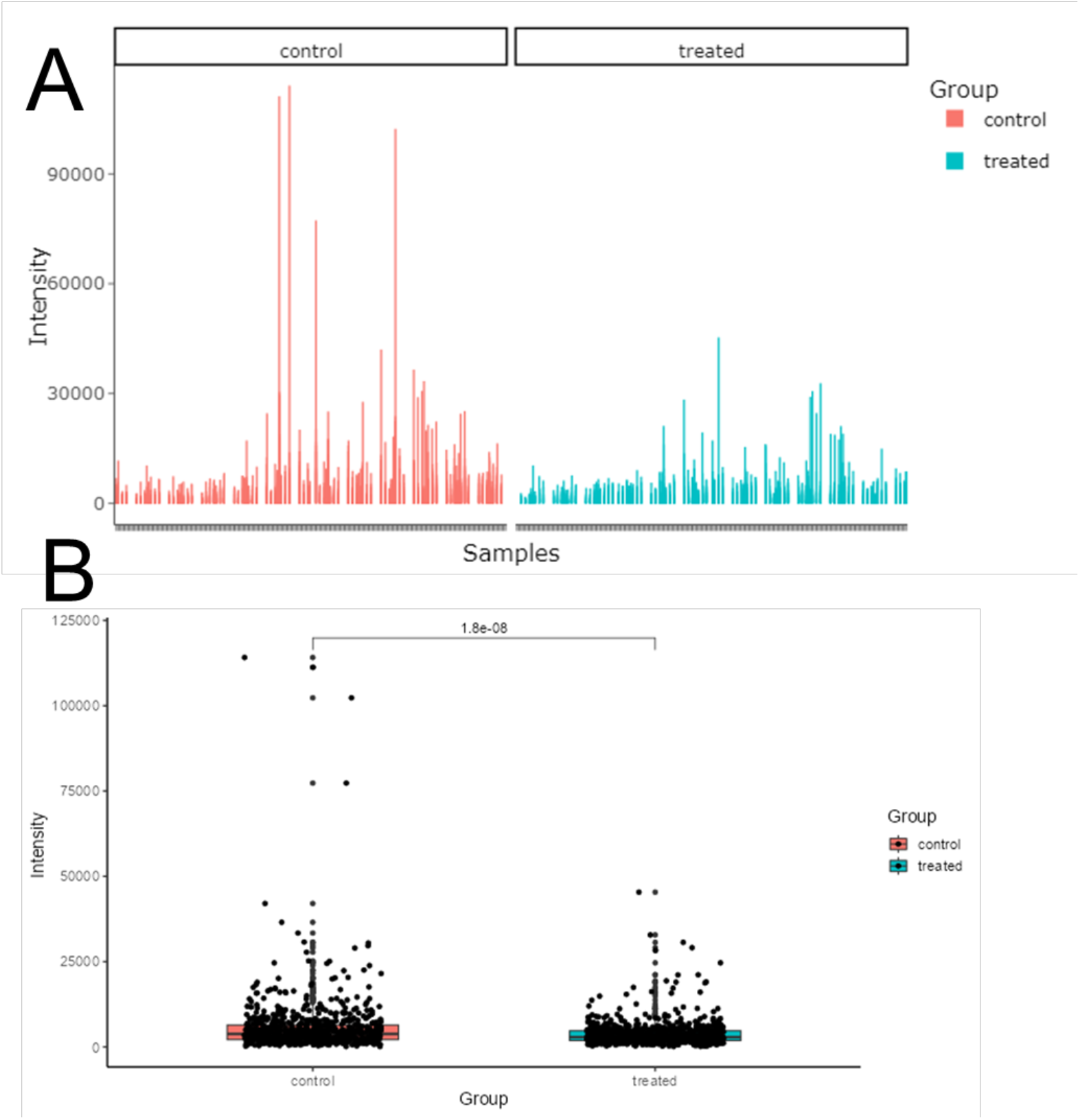
An analysis of TOP1 expression in A549 cells treated with campothecin for 48 hours. **A**. Bar plots displaying the original intensities of TOP1for each single cell analyzed at the 48 hour time point **B**. A boxplot demonstrating the significance in this alteration and the visualization of protein expression in each individual cell.

### SCP Viz allows single protein analysis across newly discovered subcellular populations

One challenge in single cell analysis is rapidly separating out cells exhibiting specific phenotypes. Orsburn *et al*., addressed this challenge by using relational databases through the GlueViz program in Anaconda Navigator.^29^ However, beyond basic visualization, extracting data from this solution requires expertise in the Python coding language. In contrast, SCP Viz provides a straightforward solution, allowing users to extract text data directly from interactable plots. SCP Viz utilizes combinatorial text filters that enable identification, labeling, and separation of new populations. In the dataset from Velgari *et al*., re-evaluated in **Figure 2**, approximately 10 control cells demonstrate markedly higher expression of TOP1 than any other cell line analyzed. These cells can be identified by using either a box select or lasso select function. Once selected in this manner, and the process executed, these cells can be downloaded as a new text file. These cells can be reloaded into SCP Viz directly for direct analysis of this subpopulation, or by relabeling these cells manually within the original report. When appending the final report this allows comparisons between control, populations with “normal” TOP1 expression, and cells with higher relative levels of expression of this protein.

### SCP Viz is compatible with single cell transcriptomics data

To further assess SCP Viz’s compatibility, we downloaded the supplementary data from a recent scSeq study where cancer cells were treated with a KRAS^G12C^ inhibitor.^30^ To simplify our analysis we focused solely on the H358 cancer cells that underwent this treatment. The input CSV file contained approximately 1,000 cells each from control, 2, 24 and 72 hours of treatment with the drug. The output data from this study already had log-transformed transcript reads, so transformation was not necessary. As shown in **Figure 3**, we had no challenge processing this entire time course study.

**Figure 3.**
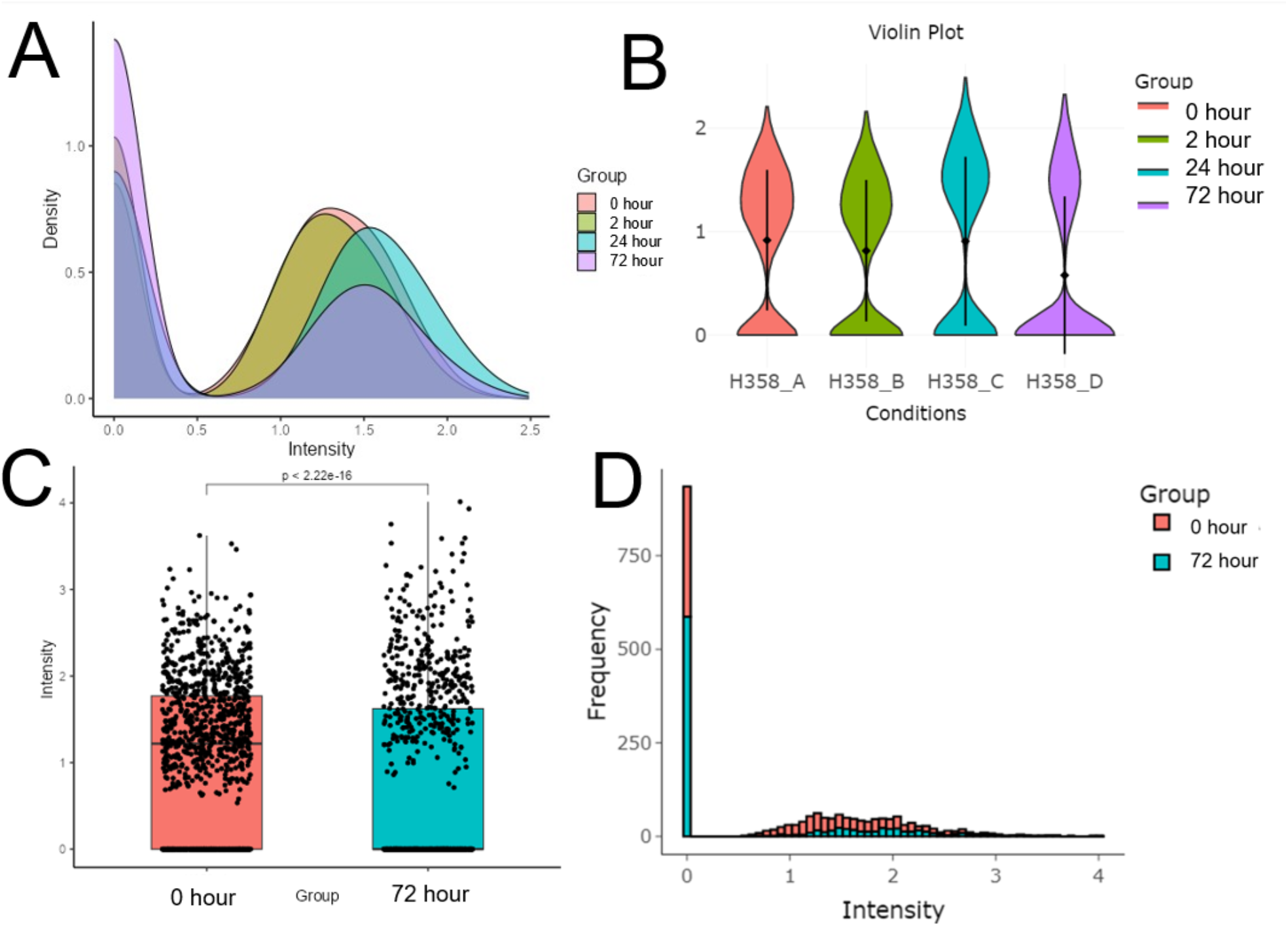
A demonstration that SCP Viz can interpret KRAS transcript expression in a scSeq analysis of a drug treatment time course of approximately 4,000 single H358 cells. **A**. A density plot representing the time course. **B**. Volcano plots displaying these same time points. **C**. A bar plot demonstrating significant downregulation of KRAS transcripts at 72 hours when compared to control cells. **D**. A histogram based on the time points from **C** demonstrates the substantial increase in zero read values for KRAS transcripts at 72 hours compared to control.

In our lab’s previous integration of these scSeq data with SCP data, we focused solely on control cells and those treated with inhibitor for 72 hours.^29^ This limitation arose because we could not effectively visualize alterations in a dataset containing 4,000 single cells. Using SCP Viz, we can now visualize the abundance of specific gene products such as KRAS across all time points in the study. With the ability to visualize the entire time course we observed a surprising increase in KRAS transcript abundance after 24 hours of treatment, relative to control. (**Figure 4A/4B**). This abundance then decreases to significantly lower levels by 72 hours, as previously observed (**Figure 4C**). From the overlapping histograms (**Figure 4D**), it is evident that there are roughly twice as many cells with undetectable KRAS transcripts at 72 hours in comparison to the control. As noted by others, one benefit of the recent advances in KRAS small molecule inhibitors is to provide new insights into the mechanisms of KRAS functionality and the complex nature of KRAS mutant driver mutations.^31,32^ Visualization of KRAS transcript abundance suggests that the earliest response to covalent inhibitors is to increase transcription rates leading to increased numbers of KRAS mRNA at 24 hours. By 72 hours, however, the number of cells producing detectable levels of KRAS mRNA is reduced by over 50% within the population. Similarly, the cells with detectable KRAS mRNA levels demonstrate a significant decrease in numbers of these transcripts.

## Conclusions

We have introduced a new tool, SCP Viz, designed for targeted analysis of single proteins across populations of single human cells. SCP Viz prioritizes compatibility, ensuring visualization regardless of the instrument and data processing software used. As further evidence of the universal nature of this tool, we have demonstrated that SCP Viz can also visualize single transcripts across a 4,000 single cell scSeq dataset. With the ability to visualize single cell responses across a time course of thousands of cells, SCP Viz allows new insights to be quickly obtained in the mechanism of action of KRAS^G12C^ covalent inhibitors.

## Acknowledgements

This work would not have been possible without suggestions and guidance of Dr. Alexis Norris. We would also like to thank Benjamin Sparklin for innovative insights leading to the keyword text filters used in SCP Viz.

